# Task dependence of neural representations of scene attributes but not scene categories in the prefrontal cortex

**DOI:** 10.1101/2020.12.04.412445

**Authors:** Yaelan Jung, Dirk B. Walther

## Abstract

Natural scenes deliver rich sensory information about the world. Decades of research has shown that the scene-selective network in the visual cortex represents various aspects of scenes. However, less is known about how such complex scene information is processed beyond the visual cortex, such as in the prefrontal cortex. It is also unknown how task context impacts the process of scene perception, modulating which scene content is represented in the brain. In this study, we investigate these questions using scene images from four natural scene categories, which also depict two types of scene attributes, temperature (warm or cold), and sound level (noisy or quiet). A group of healthy human subjects from both sexes participated in the present study using fMRI. In the study, participants viewed scene images under two different task conditions; temperature judgment and sound-level judgment. We analyzed how these scene attributes and categories are represented across the brain under these task conditions. Our findings show that scene attributes (temperature and sound-level) are only represented in the brain when they are task-relevant. However, scene categories are represented in the brain, in both the parahippocampal place area and the prefrontal cortex, regardless of task context. These findings suggest that the prefrontal cortex selectively represents scene content according to task demands, but this task selectivity depends on the types of scene content; task modulates neural representations of scene attributes but not of scene categories.

**Significance statement:** Research has shown that visual scene information is processed in scene-selective regions in the occipital and temporal cortices. Here, we ask how scene content is processed and represented beyond the visual brain, in the prefrontal cortex (PFC). We show that both scene categories and scene attributes are represented in PFC, with interesting differences in task dependency: Scene attributes are only represented in PFC when they are task-relevant, but scene categories are represented in PFC regardless of the task context. Taken together, our work shows that scene information is processed beyond the visual cortex, and scene representation in PFC reflects how adaptively our minds extract relevant information from a scene.

## Introduction

Real-world scenes provide a wealth of complex sensory experiences. Research has demonstrated that various components of scenes are processed across several brain regions of the scene-selective network in the visual cortex, such as the parahippocampal place area (PPA, Epstein & Kanwisher, 1998), the retrosplenial cortex (RSC, Maguire, 2001), and the occipital place area (OPA, Dilks et al., 2013). However, recent studies have shown that visual scene processing takes place beyond the visual cortex and involves the associative cortex such as the parietal (Silson et al., 2016; Silson et al., 2019) and the prefrontal cortex (PFC, Jung, Larson, & Walther, 2018). How are scene categories and other scene attributes represented beyond the scene-selective network? In the current study, we investigate whether and how scene content is processed beyond the visual cortex, especially in PFC.

We have previously demonstrated that scene images and sounds elicit neural representations of scene categories in PFC. Intriguingly, these representations are modality-independent, suggesting that PFC represents scene content in an abstract way, not constrained in a single sensory modality (Jung et al., 2018). Still, we do not know yet what elements of scene content are processed and represented in PFC. That is, natural scenes also comprise other scene attributes than just scene categories, some (those related to spatial context) often referred to as global scene properties, such as openness, navigability, or transience (Greene & Oliva, 2009), which are also processed in scene-selective visual brain regions (Kravitz et al., 2011; Park et al., 2011; Persichetti & Dilks, 2019). However, we do not know yet whether the PFC represents scene attributes that are not related to categories, as it does scene categories, and whether the neural presentations of scene attributes in PFC are also modality-independent.

Scene representations in PFC may also depend on task context. One of the PFC’s fundamental functions is to modulate the neural representation of sensory inputs according to task demands (Chadick & Gazzaley, 2011; Harel et al., 2014; Zanto et al., 2011). Thus, although scene content can be represented in PFC, it could be only when such scene content is task-relevant. Specifically, we hypothesize that task context is more likely to impact the processing of scene attributes than that of scene categories, as supported by previous research. First, behavioral studies have shown that scene category information is accessible with limited attention (Li et al., 2002), and even when it is not relevant to the task context (Greene & Fei-Fei, 2014). Second, it has been shown that scene categories can be represented in PFC in a passive-viewing paradigm (Jung et al., 2018; Walther et al., 2009), which does not explicitly require attention to scene categories. Thus, we predict that scene category information may be represented in PFC regardless of task context. We explore these possibilities by assessing the neural representations of scene attributes and scene categories in PFC under different task conditions.

The current study employs two scene attributes (temperature and sound-level) and four natural scene categories (beaches, cities, forest, and train stations) to examine how scene attributes and scene category information are represented in PFC across different task contexts. To construct different task conditions for each scene property, we instructed participants to make a judgment on a scene attribute, which would encourage them to attend to a particular property in a scene.

Finally, the present study also explores whether representations of scene attributes in PFC are modality-independent in the same way as scene categories (Jung et al., 2018). We do so by comparing the neural representation of scene attributes (both temperature and sound-level) inferred from images to the neural representation of direct thermal and auditory percepts. Our findings demonstrate that both scene attributes and scene categories are represented in PFC in a modality-independent way. Neural representations of scene attributes generalize across sensory modalities, as indicated by similar neural activity patterns across percepts inferred from images and direct sensation.

## Materials & Methods

### 1. Participants

Twenty-five healthy adults recruited from the University of Toronto participated in the fMRI study (10 male, 15 females; mean age = 22.17, SD = 1.99, all right-handed). All participants reported normal or corrected-to-normal vision and normal hearing and no history of neurological abnormalities. Five participants did not complete both sessions, and their data were excluded from the analysis. Participants provided informed consent before the experiment, and all aspects of the experiment were approved by the Research Ethics Board (REB) at University of Toronto (protocol number 33720).

### 2. Experiment design & Stimuli

The study consisted of two sessions, the main experiment session and the localizer session. For all participants, the main experiment session was performed first, except for three, who did the localizer experiment first for logistical reasons (e.g., last-minute cancellation). The main experiment session consisted of three different types of tasks: fixation monitoring, temperature judgment, and sound-level judgment. Participants always performed two runs of the fixation task first, and then the other two tasks in alternation (either T-S-T-S or S-T-S-T). The localizer session consisted of three different experiments; the sound experiment, the stone experiment, and the face-place-object localizer, which were always performed in this order.

#### 2-1. Visual Stimuli

Sixty-four scene images from four natural scene categories were selected according to three scene properties: entry-level category (beaches, city streets, forests, and train stations), temperature of the depicted scene (warm versus cold), and sound level (noisy versus quiet). The selection procedure ensured that all three properties vary *independently* within the image set.

We measured the properties of scene images with a set of three behavioral experiments on Amazon MTurk. Workers in the behavioral experiments viewed each image and rated the temperature and the sound level on a seven-point Likert scale. Each type of property was measured in a separate block. The order of the tests was counterbalanced across participants. Before the rating, a detailed description of each scale was provided, followed by six practice trials separately for the temperature rating and the sound-level rating (see Supplementary Figure 1).

In the first rating experiment, 27 MTurk workers rated 292 scene images which were collected from the internet. Ratings were z-scored for each participant and averaged across participants for each image. We excluded 87 images with extreme temperature or sound scores, added 100 new images, and repeated the rating experiment with 15 participants, first on 305 images and then on a subset 128 images. After this procedure, we selected the final set of 64 images and repeated the rating experiment on these images with 45 participants. These final ratings show that neither temperature nor sound level are related to category labels. A Kruskal-Wallis test was conducted to find the effect of category labels on the temperature rating, showing that there is no effect of category labels, *Kruskal-Wallis chi-square* = 3.54, *p* = 0.32; a Kruskal-Wallis test to measure the effect of category labels on sound-level ratings also reveal no effect of category labels, *Kruskal-Wallis chi-square* = 4.56, *p* = .21). Furthermore, the temperature ratings and the sound-level ratings were not correlated to each other, *r* = .037, *p* = .781 (see Figure 1D; also see Supplementary Figure 2 for the example stimuli). This final set of 64 images was used in the main experiment. The entire stimuli set and the rating data are available in a GitHub repository (https://github.com/yaelanjung/scenes_fmri).

**Figure 1.**
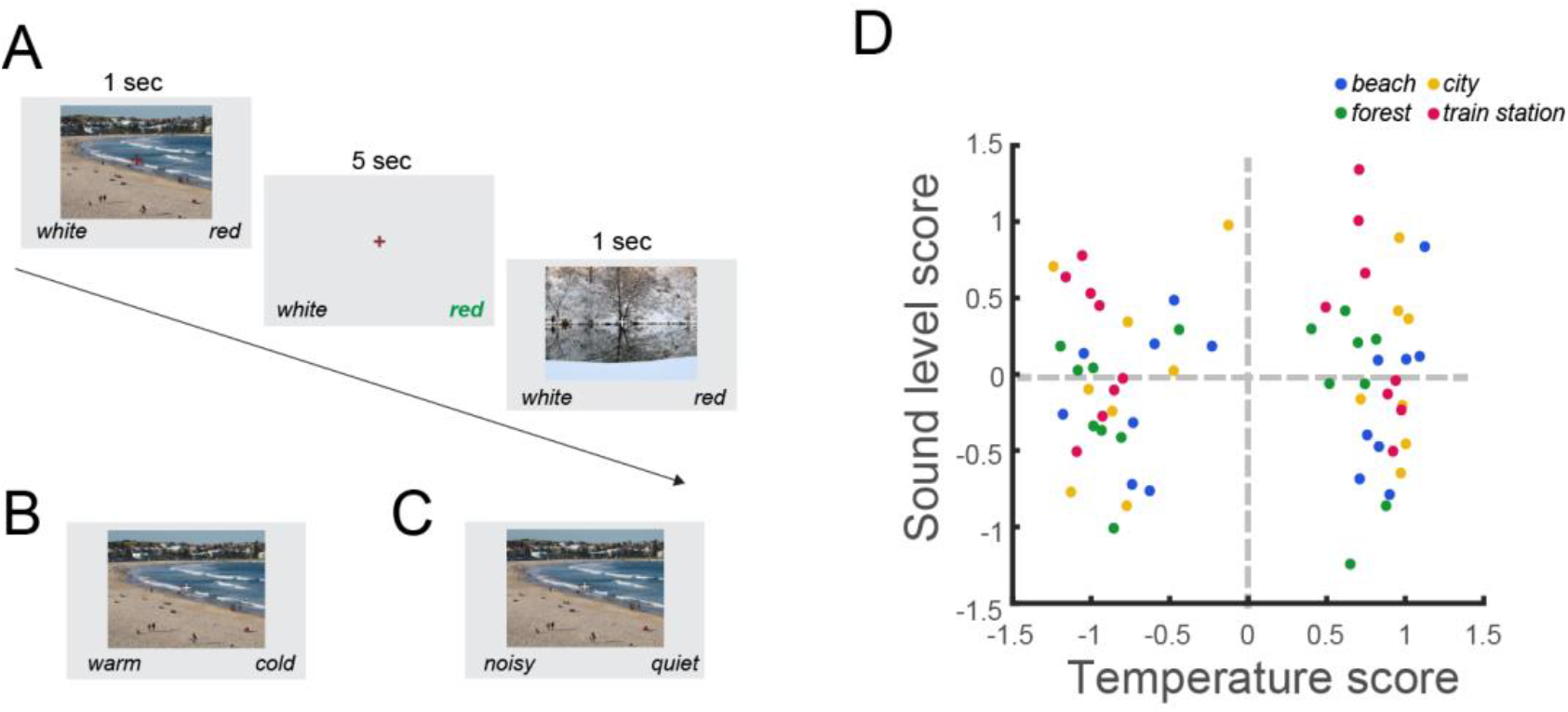
A) Illustration of the image presentation in the main experiment. B & C) Image presentation with different labels in the temperature judgment condition (B) and in the sound-level judgment condition (C). D) Behavioral ratings of temperature and sound level properties

#### 2-2. Auditory Stimuli

Ten different sounds from different places (e.g., beaches, forests, and city streets) were downloaded from an open database, Freesound (freesound.org). The sounds were cut to 12 seconds length and presented using Sensimetrics S14 MR-compatible in-ear noise-canceling headphones. The noisy and quiet sounds differed in two aspects. First, the noisy sounds had more auditory objects (i.e., birds, people, waves at a beach) than the quiet sounds. Also, based on the perceived loudness which was estimated using a sound meter over the headphones, the noisy sounds had greater volume (on average 81 dB) than the quiet sounds (on average 53 dB).

#### 2-3. Stone Stimuli

Six massage stones (approximately 6×8×2 cm) were used in the temperature experiment. To deliver stones at a particular temperature, three stones were soaked in warm water (approximately 45 °C) in a plastic thermal cup, and the other three stones were soaked in cold water (approximately 9°C) for about 10 minutes before the experiment. We measured the temperature of the stones before and after each run with a digital infrared thermometer. On average across different runs and participants, the cold stones were 11.0 °C (SD = 1.3°C) before the runs and 15.3 °C (SD = 0.93 °C) after the runs. The warm stones were 44.4 °C (SD = 0.93°C) before and 39.8 °C (0.81 °C) after the runs (see Figure 3B).

### 3. Procedure

#### 3.1. Main experiment

The main fMRI experiment was event-related with six runs of image presentations with three different tasks (two runs for each task). In all three task conditions, participants viewed the same set of 64 images one at a time for one second, with a 5-seconds inter-trial interval (Figure 1A). The trial sequence also contained 16 blank trials with only the fixation cross visible; the first and last two trials in each run were always blank, and the other twelve blank trials were randomly distributed throughout the run. Visual stimuli were presented on the screen at a distance of approximately 134 cm from the participant with images subtending approximately 12.3 degrees of visual angle. Stimulus presentation, timing of all stimuli, and recording of responses were controlled with custom Matlab scripts, using the Psychophysics Toolbox (Brainard, 1997).

In the fixation monitoring runs, participants were asked to press a button to report whether the fixation cross was white (75% of the time) or red (25% of the time). In the temperature-judgement runs, participants were asked to indicate whether the depicted scene looked warm or cold (Figure 1B). In the sound-level judgement runs, participants were asked to indicate whether the depicted scene looked noisy or quiet (Figure 1C). It was emphasized that there is no right or wrong answer in the temperature and sound-level judgement tasks. The same set of 64 images were repeated in all six runs, and the order of images was randomized in each run. Before the onset of each run, participants were reminded of the task by text on the screen as well as verbal instructions from the experimenter (behavioral data reported in Supplementary Figure 6). All participants first performed the two fixation-monitoring runs and then the temperature runs and the sound-level runs interleaved (either T-S-T-S or S-T-S-T). The order was counter-balanced across participants.

#### 3.2. Localizer experiments

The localizer experiment consisted of the sound experiment, the temperature experiment, and the face-place-object localizer. All of these experiments used a block design.

The sound experiment consisted of two runs. Each run had 10 blocks, during which 12-second sound clips were played. The clips in half of the blocks were noisy sounds, and the other half of the blocks contained quiet sounds. The order of the sound sequences was randomized for each run and for each participant. A fixation cross was presented at the center of the screen during the entire experiment.

The stone experiment consisted of four runs and was performed by two experimenters, one in the control room and one in the exam room next to the MRI scanner. Each run had six blocks, and each block was one stone event. At every onset or offset of a block, the experimenter in the control room signaled the event by presenting a colored flag (red: for warm stone, blue: for cold stone, green: for taking the stone away) through the control room window. The experimenter in the exam room dried a warm or cold stone on a towel and handed it to the left hand of the participant. Participants were holding the stone without any particular task for 10 seconds, and then the experimenter in the exam room took the stone from the participant’s hand at the green flag signal.

The face-place-object localizer experiment consisted of two runs. Each run had 4 blocks, and there were four mini-blocks with images of faces, objects, places, and scrambled object pictures in each block (Epstein & Kanwisher, 1998). Each mini block was 16 seconds long, and each image was presented for 1 second with an inter-trial interval of 500 ms. There were 12 seconds fixation periods between blocks and also at the beginning and the end of the run.

### 5. fMRI data acquisition

Imaging data were recorded on a 3 Tesla Siemens Prisma MRI scanner with a 32-channel head coil at the Toronto Neuroimaging Centre (ToNI) at the University of Toronto. High resolution anatomical images were acquired with a MPRAGE (magnetization-prepared rapid acquisition with gradient echo) protocol with multiband factor of 2. Images were then reconstructed using GRAPPA (Griswold et al., 2002), with sagittal slices covering the whole brain; inversion time TI = 912ms, repetition time TR = 1900ms, echo time TE = 2.67ms, flip angle = 9°, voxel size = 1 x 1 x 1 mm, matrix size = 224 x 256 x 160 mm. Functional images for the main and localizer experiments were recorded with a multiband acquisition sequence; TR = 2 s, TE = 32 ms, flip angle = 70 degrees, multiband factor = 4, voxel size = 2 x 2 x 2 mm, matrix size = 220 x 220 x 136 mm, 68 axial slices.

### 6. Data analyses

#### 6.1. fMRI data preprocessing

FMRI data from the main experiment runs were motion corrected to one EPI image (the middle volume in the middle run), followed by spatial smoothing with a Gaussian kernel with 2 mm full width at half maximum (FWHM). Data were converted to percent signal change with respect to the mean of each run. Preprocessing was performed using AFNI (Cox, 1996).

#### 6.2 Region of Interest

PPA was defined using data from the face-place-object localizer runs. FMRI data from this localizer were pre-processed the same way as the main experiment data, but spatially smoothed with a 4 mm FWHM Gaussian filter. Data were further processed using a general linear model (3dDeconvolve in AFNI) with regressors for all four image types (scenes, faces, objects, scrambled objects). PPA was defined as contiguous clusters of voxels with significant contrast (*q* < 0.05, corrected using false discovery rate) of scenes > (faces and objects) (Epstein & Kanwisher, 1998).

The representation of the left hand in somatosensory cortex (S1) was also functionally defined using data from the temperature localizer runs as we are interested in a region that specifically corresponds to thermal perception driven by the stone stimuli. fMRI data from the temperature localizer runs were pre-processed the same way as the main experiment data and spatially smoothed with a 4mm FWHM Gaussian Filter. Data were then further processed using a general linear model (3dDeconvolve in Afni) with regressors for stone events (stone, and no stone), irrespective of temperature. S1 for the left hand was defined as a contiguous cluster of voxels in the right hemisphere with significant contrast (*q* < 0.05, corrected using false discovery rate) of stone > no-stone. In all participants except one, we observed a cluster with significant contrast near the central gyrus in the right hemisphere (contralateral to the left hand where the stone was given). The cluster was then masked by the anatomical extent of the post-central gyrus, which is known as the location of the primary somatosensory cortex (see Supplementary Figure 3). The subject for whom we could not functionally define S1 was excluded from the decoding analysis performed in this area.

Anatomically defined ROIs were extracted using a probabilistic atlas in AFNI (DD Desai MPM; Destrieux et al., 2010): middle frontal gyrus (MFG), superior frontal gyrus (SFG), and inferior frontal gyrus (IFG). Anatomical masks for primary auditory cortex (Heschl’s gyrus) were made available by Sam Norman-Haignere (Norman-Haignere et al., 2013). After nonlinear alignment of each participants’ anatomical image to MNI space using AFNI’s 3dQwarp function, the inverse of the alignment was used to project anatomical ROI masks back into original subject space using 3dNwarpApply. All decoding analyses, including for the anatomically defined ROIs, were performed in original subject space.

#### 6.3. Decoding analyses

##### 6.3.1. Multi-voxel pattern analysis of the main experiment

Decoding analysis was performed for each task condition, the temperature judgment condition and the sound-level judgment condition. Data from the fixation-monitoring condition were included as training but not as test data for both conditions. For each participant, we trained a linear support vector machine (SVM; using LIBSVM, Chang & Lin, 2011) to assign the correct labels to the voxel activations inside an ROI based on the fMRI data from three runs: one of the task-specific runs as well both of the fixation-monitoring runs. The SVM decoder then produced predictions for the labels of the data in the left-out task run. This cross-validation procedure was repeated with the other task run left out, thus producing predicted labels for both task-specific runs. This procedure was performed for scene categories (beaches, forests, city streets, train stations), scene temperature (warm, cold), and sound level (noisy, quiet) separately for temperature judgment and sound-level judgement runs. Decoding accuracy was assessed as the fraction of predictions with correct labels. Group-level statistics was computed over all twenty participants using one-tailed t tests, determining if decoding accuracy was significantly above chance level (0.5 for temperature and sound-level and 0.25 for scene categories). In all the statistical tests performed here, we adjusted significance of the t-test for multiple comparisons using false discovery rate (FDR) (Westfall & Young, 1993).

To curb over-fitting of the classifier to the training data, we reduced the dimensionality of the neural data by selecting a subset of voxels in each ROI. Voxel selection was performed by ranking voxels in the training data according to the F statistics of a one-way ANOVA of each voxel’s activity with temperature, sound level, or scene categories as the main factor (Pereira et al., 2009). We determined the optimal number of voxels by performing a nested leave-one-run-out (LORO) cross validation within the training data. It should be noted that the voxel selection is performed only using the training data (nested cross-validation) to avoid double-dipping. Once an optimal number of voxels was determined by voxel selection, a classifier was trained again using the entire training data (but with selected voxels) and tested on the test data. The test data were not used to select voxels or train the classifier for a given cross-validation fold. Optimal voxel numbers varied by ROI and participant, showing an overall average of 99 voxels across all ROIs and participants in the analysis of the main experiment.

##### 6.3.2. Multivariate pattern analysis of the localizer experiments

To decode temperature from the localizer experiment, we performed a GLM analysis on the preprocessed data with regressors for warm and hot blocks as well as nuisance regressors for motion and scanner drift within the run. We obtained eight sets of beta weights (4 runs x 2 conditions) and performed a LORO cross-validation analysis using an SVM classifier. A voxel selection procedure (as described above) was also included in this decoding procedure. For each cross-validation fold, the voxels were ranked based on the F statistics of an ANOVA of each voxel’s activity with temperature as the factor. We determined the optimal number of voxels by performing a LORO cross-validation analysis within the training data (nested cross-validation; mean = 211.11).

There were only two runs of data in the sound experiment. Thus, we performed a GLM analysis at the level of blocks instead of runs. We used the sets of beta weights for each block as input to the decoder. Specifically, in this GLM analysis, the preprocessed data from the sound localizer runs were analyzed with regressors for noisy and quiet sound blocks as well as nuisance regressors for motion and scanner drift within the run (second-order polynomial). As a result, we obtained twenty sets of beta weights (10 blocks x 2 conditions). These sets of betas were used in a one-block-leave-out cross-validation analysis (similar to LORO, but on each block instead of runs) using an SVM. A voxel selection procedure (as described above) was also performed for each fold in the decoding procedure, and average optimal number of voxels (across all ROIs) was 304.02. The decoding accuracy for each ROI and each subject was determined by averaging the performance across all the folds of the cross-validation procedure.

##### 6.3.3. Cross-decoding analysis

In the cross-decoding analysis, we trained a classifier on data from the localizer session and tested it with data from the main experiment. Since training and test data were completely separate in these conditions, we used all data from the localizer session for training and voxel selection. We here used the accuracy values from the whole-brain searchlight analysis to rank voxels (see the following section for the detail). Using the training data, we performed LORO cross validation analyses with the number of selected voxels varying from 125 to 500 (step size of 25). We included voxels according to decreasing rank order of their decoding accuracies in the searchlight analysis. We compared the decoding performance for different voxel numbers and determined the optimal number of voxels (mean = 171.25), which showed the best decoding performance within the training data. The decoder was then trained on the entire training set, using the optimal number of voxels, and then tested on the data from the main experiment at those same voxel locations.

##### 6.3.4. Searchlight analysis

To explore representations of scene categories outside of pre-defined ROIs, we performed a searchlight analysis. We defined a cubic “searchlight” of 5×5×5 voxels (10×10×10 mm). The searchlight was centered on each voxel in turn (Kriegeskorte et al., 2006), and LORO cross-validation analysis was performed within each searchlight location using a linear SVM classifier (CosmoMVPA Toolbox; Oosterhof et al., 2016). Decoding accuracy at a given searchlight location was assigned to the central voxel.

For group analysis, we first co-registered each participant’s anatomical brain to the Montreal Neurological Institute (MNI) 152 template using a diffeomorphic transformation as calculated by AFNI’s 3dQWarp. We then used the same transformation parameters to register individual decoding accuracy maps to MNI space using 3dNWarpApply, followed by spatial smoothing with a 4 mm FWHM Gaussian filter. To identify voxels with decodable categorical information at the group level, we performed one-tailed t-tests to test whether decoding accuracy at each searchlight location was above chance (0.5 for non-visual properties and cross-decoding, and 0.25 for categories). After thresholding at *p* < 0.05 (one-tailed) from the t-test, we conducted a cluster-level correction for multiple comparisons. We used AFNI’s 3dClustSim to conduct alpha probability simulations for each participant. The estimated smoothness parameters computed by 3dFWHMx were used to conduct the cluster simulation. In the simulations, a p-value of 0.05 was used to threshold the simulated data prior to clustering and a corrected alpha of 0.05 was used to determine the minimum cluster size. We used the average of the minimum cluster sizes across all the participants as the cluster threshold, which was 215 voxels.

## Results

### Decoding of scene attributes

To assess neural representations of temperature and sound level from images, we performed the multivariate pattern analysis (MVPA) for each ROI and for each task condition.

As illustrated in Figure 2A, we were able to decode temperature (warm versus cold) in the PPA, mean decoding accuracy = 58.75%, *t*(19) = 2.67, *q* = 0.01, and in MFG, mean decoding accuracy = 62.5%, *t*(19) = 2.94, *q* = 0.02, when participants performed the temperature judgment task. However, when participants were performing the sound-level judgment task, the temperature property was decoded neither in the PPA, mean accuracy = 50%, *t*(19) = 0, *q* = 0.62; nor in the MFG, mean accuracy = 50%, *t*(19) = 0, *q* = 0.62 (see Table 1, for decoding performance in other ROIs). Sound level (noisy versus quiet) was decoded in the sound-level judgment condition, in the MFG, mean accuracy = 62.5%, *t*(19) = 2.52, *q* = 0.01 and the IFG, mean accuracy = 63.75%, *t*(19) = 2.98, *q* = 0.02. However, when the participants were performing the temperature judgment task, sound level could not be decoded in these areas, in MFG, mean accuracy = 55%, *t*(19) = 0.94, *q* = 0.18; in IFG, mean accuracy = 48.75%, *t*(19) = −0.29, *q* = 0.61.

**Figure 2.**
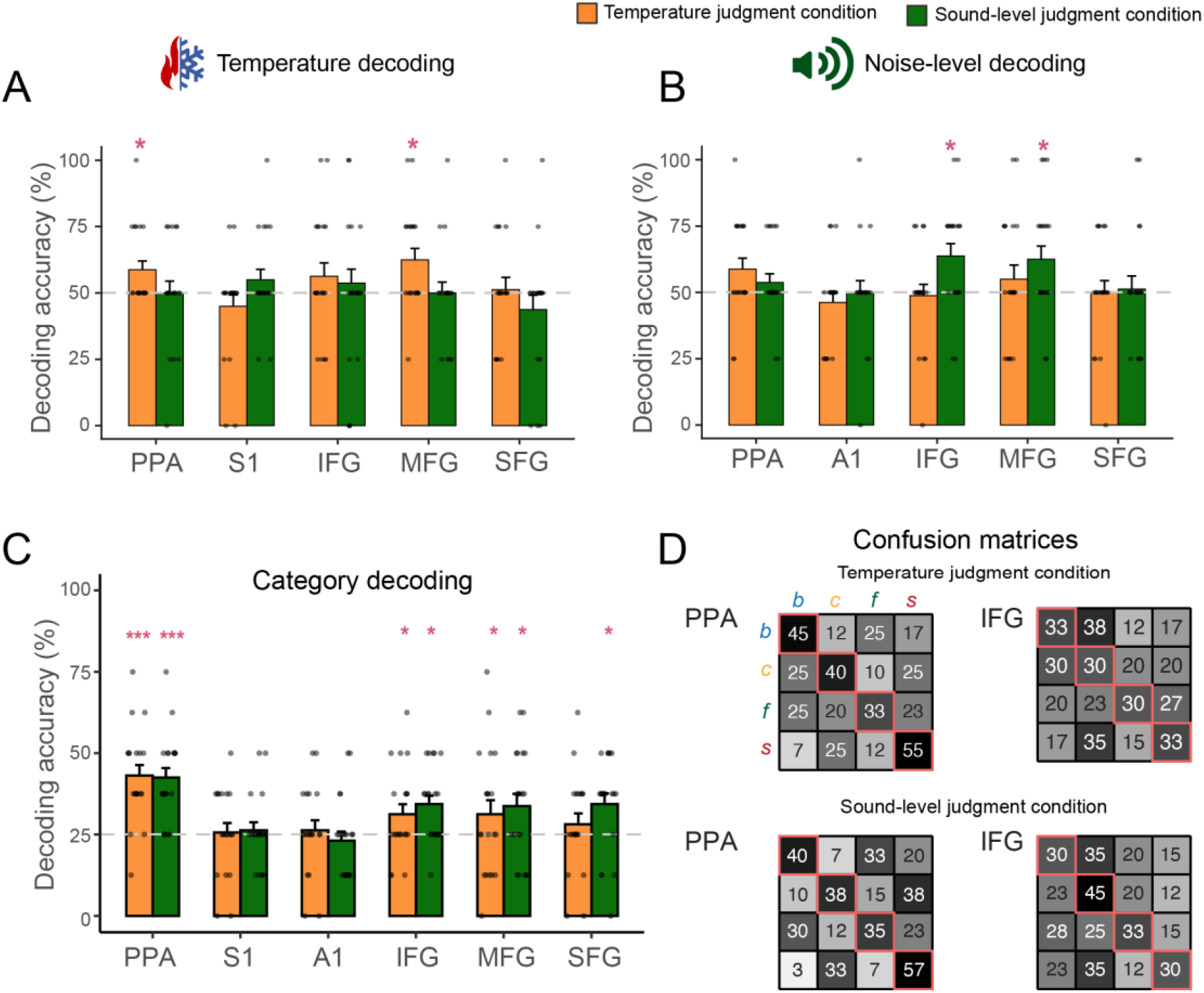
Decoding from the main experiment. Decoding of temperature (A), sound-level properties (B), and categories (C) of scenes in each ROIs in the temperature judgment condition (orange bars) and in the sound-level judgment condition (green bars). (D). Confusion matrices from category decoding: each row indicates presented category (in order of beach, city, forest, and station), and each column indicates the predicted category. Diagonal boxes (with a pink frame) indicate percent of corrected predictions (rounded to the nearest integer). Dots in each figure represent individual data points. * *q*< 0.05, *** *q* < 0.001

**Table 1.** Mean of decoding accuracy and statistical analysis in each ROI for the temperature and the sound-level properties

| ROI | Decoding of temperature |  |  |  |  |  |  |  | Decoding of sound-level |  |  |  |  |  |  |  |
| --- | --- | --- | --- | --- | --- | --- | --- | --- | --- | --- | --- | --- | --- | --- | --- | --- |
|  | Temperature condition |  |  |  | Sound-level condition |  |  |  | Temperature condition |  |  |  | Sound-level condition |  |  |  |
|  | mean | t stats | q | effect size | mean | t stats | q | effect size | mean | t stats | q | effect size | mean | t stats | q | effect size |
| PPA | 58.75 | 2.67 | 0.01 | 0.60 | 50.00 | 0.00 | 0.50 | 0.00 | 58.75 | 2.10 | 0.12 | 0.47 | 53.75 | 1.14 | 0.22 | 0.26 |
| S1 | 50.00 | 0.00 | 0.50 | 0.00 | 52.63 | 0.81 | 0.21 | 0.18 | - | - | - | - | - | - | - | - |
| A1 | - | - | - | - | - | - | - | - | 46.25 | -1.00 | 0.84 | -0.22 | 50.00 | 0.00 | 0.50 | 0.00 |
| MFG | 62.50 | 2.94 | 0.02 | 0.66 | 50.00 | 0.00 | 0.50 | 0.00 | 55.00 | 0.94 | 0.45 | 0.21 | 62.50 | 2.52 | 0.03 | 0.56 |
| SFG | 51.25 | 0.27 | 0.49 | 0.06 | 43.75 | -1.16 | 0.87 | -0.26 | 50.00 | 0.00 | 0.77 | 0.00 | 51.25 | 0.25 | 0.50 | 0.06 |
| IFG | 56.25 | 1.23 | 0.20 | 0.27 | 53.75 | 0.72 | 0.60 | 0.16 | 48.75 | -0.29 | 0.77 | -0.07 | 63.75 | 2.98 | 0.02 | 0.67 |

For scene temperature, it is possible that the color of the images may hold cues to the temperature. That is, the classifier might be mainly relying on color information, not necessarily perceived scene temperature information. We explored this possibility by controlling for the color information in the fMRI data. We calculated the average L, a, b values from the CIE-LAB color channels for individual images and regressed out all of these values from the fMRI data before we perform the GLM analysis. When we performed the MVPA on the color-controlled betas from the GLM analysis, we found that decoding accuracy is impacted in the PPA, mean accuracy = 53.75%, *t*(19) = 1.14, *p* = 0.1, but not in the MFG, mean accuracy = 58.75%, *t*(19) = 2.10, *p* = 0.02. Thus, it appears that the neural representation of scene temperature in the PPA but not the MFG strongly relies on color information (also see Supplementary Figure 4).

To directly examine the effect of task context on decoding of scene attributes, we performed a two-way ANOVA for each ROI with task (temperature-judgment versus sound-level judgment) and decoded property (temperature versus sound level) as a factor. In all ROIs, there was no main effect of either task or decoded property [all *Fs* < 3.32, *ps* > .07], as well as no significant interaction between the two [all *Fs* < 3.3, *ps* > 0.07, except in MFG. In the MFG, although there was no main effect of task, or decoded property, the interaction between decoded property and task was significant, *F*(1,76) = 4.57, *p* = 0.003. These results indicate that task context impacts which type of non-visual properties are represented in the MFG.

### Decoding of scene categories

Scene categories were decoded with a linear SVM classifier using the same data from each task condition but with category labels as ground truth. Here, chance level was 25% since there were four different scene categories.

As illustrated in Figure 2C, scene categories were decoded in both task conditions in PFC as well as in PPA. In the temperature judgement condition, decoding accuracies for scene categories were significantly higher than chance in PPA, mean accuracy = 42.50%, *t*(19) = 5.98, *q* < 0.001; in MFG, mean accuracy = 33.75%, *t*(19) = 2.33, *q* = 0.01; and in IFG, mean accuracy = 34.38%, *t*(19) = 3.68, *q* = 0.02. In the sound-level judgment condition, decoding accuracy for scene categories was significantly higher than chance in PPA; mean accuracy = 43.14%, *t*(19) = 5.78, *q* < 0.001; in IFG, mean accuracy = 31.25%, *t*(19) = 2.03, q = 0.04, in MFG, mean accuracy = 33.75%, *t*(19) = 2.33, *q* = 0.03, and in SFG; mean accuracy = 34.38%, *t*(19) = 2.88, *q* = 0.01 (see Table 2 for other ROIs). The confusion matrices from decoding performance in each ROI show that the classifiers successfully decode all four categories (Figure 2D; Supplementary Figure 5). To test whether the task impacts decoding accuracy of scene categories, one-way ANOVA was performed on decoding accuracy with task as a factor in each ROI. In all ROIs, there was no significant effect of task on decoding accuracy for scene categories, all *Fs* <1.55, all *ps* > 0.22. Thus, these findings indicate that unlike scene attributes, scene categories are represented in PFC regions regardless of task context.

**Table 2.** Mean of decoding accuracy and statistical analysis in each ROI for scene category

| ROI | Decoding of scene category |  |  |  |  |  |  |  |
| --- | --- | --- | --- | --- | --- | --- | --- | --- |
|  | Temperature judgment task |  |  |  | Sound-level judgment task |  |  |  |
|  | mean | t stats | q | effect size | mean | t stats | q | effect size |
| PPA | 43.13 | 5.66 | < .001 | 1.27 | 42.50 | 5.98 | < .001 | 1.34 |
| S1 | 25.63 | 0.21 | 0.74 | 0.04 | 26.32 | 0.52 | .36 | 0.12 |
| A1 | 26.32 | 0.42 | 0.34 | 0.09 | 23.13 | -0.68 | 0.75 | 0.09 |
| MFG | 33.75 | 2.33 | 0.01 | 0.33 | 33.75 | 2.33 | 0.02 | 0.52 |
| SFG | 28.13 | 1.56 | 0.23 | 0.20 | 34.38 | 2.88 | 0.01 | 0.64 |
| IFG | 31.25 | 2.03 | 0.04 | 0.45 | 34.38 | 3.68 | 0.002 | 0.82 |

As we identified the neural representations of temperature and sound level in scene images, we here further explore these representations to better understand how scene content is represented in PFC. Previous research has demonstrated that the prefrontal cortex represents scene categories at an abstract-level, which generalizes between visual and auditory input (Jung et al., 2018). Thus, it is possible that the prefrontal cortex represents scene attributes, temperature and sound-level information, at such an abstract level as well. To examine this possibility, we first tested how both properties are represented in the brain when participants perceived them through direct sensory stimulation. We then performed cross-decoding analyses from direct sensations to percepts inferred from images to explore whether scene attributes are also represented beyond specific sensory domains.

### Decoding of direct percepts & cross-decoding: temperature properties

We first examined the neural representation of temperature that arises from direct thermal sensation. In the localizer experiment, participants were holding a stone in their left hand which was either warm (around 42°C, averaged across before and after each run) or cold (around 13 °C, averaged across before and after each run) (see Figure3A & B). We performed a LORO cross-validation analysis on these data to decode the temperature across different brain regions. In this analysis, we included the representation of the left hand in somatosensory cortex. Since the stone stimulus was always handed to the left hand of participants, we only considered the contralateral (right) side of somatosensory cortex. We also included the prefrontal regions, inferior, middle, and superior frontal gyri, in which we observed neural coding of inferred thermal sensation from the main experiment.

**Figure 3.**
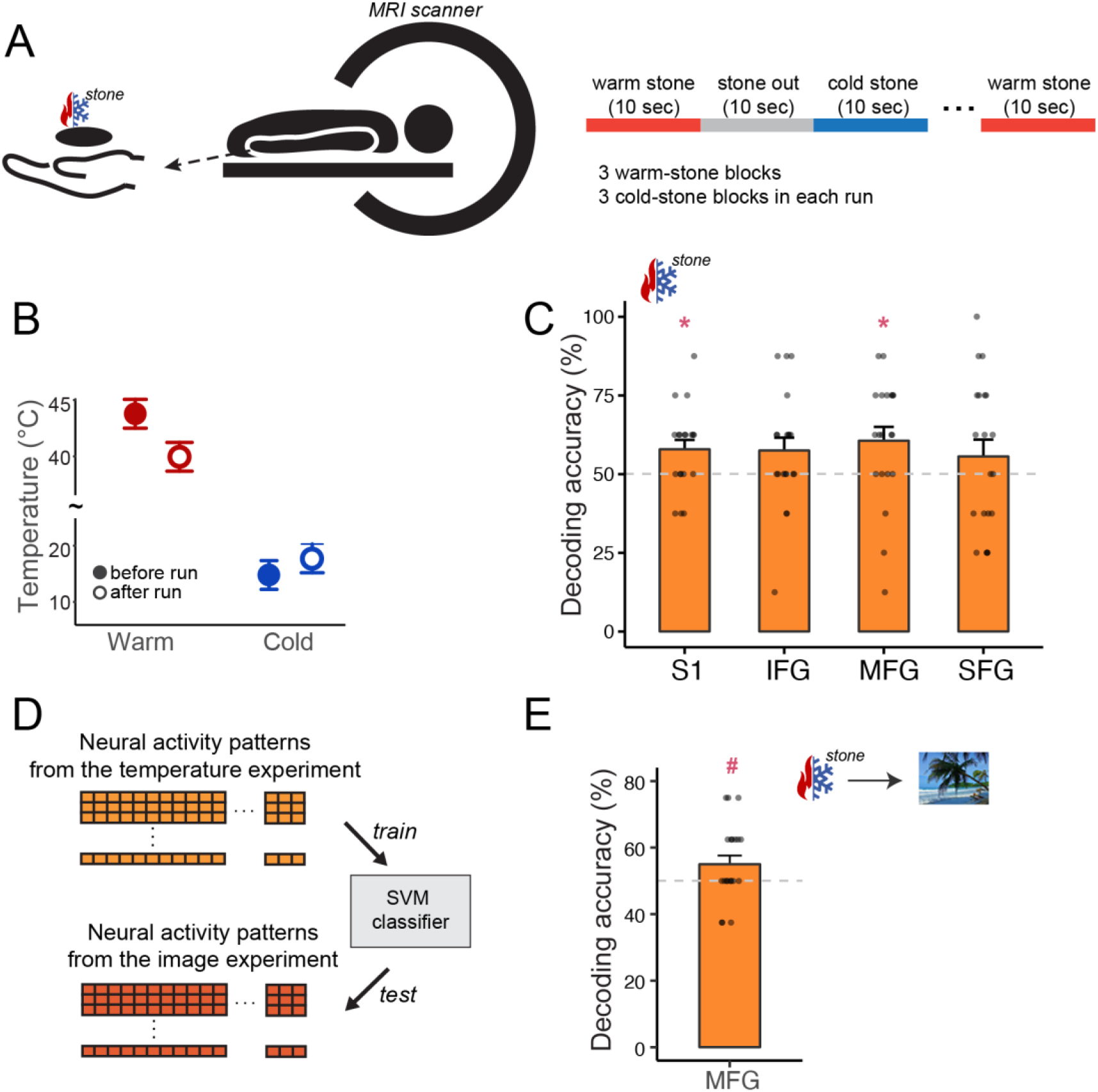
Decoding of temperature property. A) Illustration of the stimuli presentation and the experiment design for the temperature localizer runs. B) Temperature of warm and cold stones before and after runs. C) Decoding of temperature from thermal sensations. D) Schematic of cross-decoding analysis across direct and indirect thermal percepts. E) Decoding accuracy of the cross-decoding analysis in the post-central sulcus and MFG. Dots in each figure represent individual data points. * *q* < 0.05, # *p* < 0.05

As illustrated in Figure 3C, the decoder was significantly more accurate than chance (50%) in the right S1, mean accuracy = 57.89%, *t*(18) = 2.584, *q* = 0.031, suggesting that the temperature of the stones is represented in S1. Furthermore, the classifier can predict the temperature of the stones in the MFG, mean accuracy = 60.62%, *t*(19) = 2.43, *q* = 0.044, indicating that, just like inferred temperature information from scene images, the thermal percept is also represented in PFC.

Having identified brain regions that represent temperature delivered from direct sensations, we here investigate whether such neural representation is similar to the percepts inferred from images, which we observed in the main experiment. We do so by performing a cross-decoding analysis in the ROIs that showed significant decoding of sensory properties from images as well as direct percepts. For the temperature property, only MFG shows significant decoding of temperature from images (Figure 2A) and from direct percepts (Figure 3C). An SVM classifier was trained on the data from the direct percept of temperature in the localizer experiment and then tested on the data from viewing scene images in the main experiment (Figure 3D).

Cross-decoding from the stone experiment to the main scene image experiment was significantly above chance in MFG, mean accuracy = 55%, *t*(19) = 1.9, *p* = 0.036, suggesting that the representation of temperature generalizes well from direct to inferred percepts in this brain region (Figure 3E).

### Decoding of direct percepts & cross-decoding: sound-level properties

Similarly, we examined whether direct auditory sensations generalize to the neural representations of inferred sounds which were identified in the main experiment. First, we tested neural representations of noisy and quiet sounds from actual auditory percepts; an SVM classifier was trained and tested on the neural activity patterns from the sound localizer runs using LORO cross-validation. As illustrated in Figure 4B, the decoder correctly classified the sound level of scene sounds in A1, mean accuracy = 84%, *t*(19) = 9.95, *q* < 0.001, and in the IFG, mean accuracy = 58%, *t*(19) = 3.046, *q* = 0.006.

**Figure 4.**
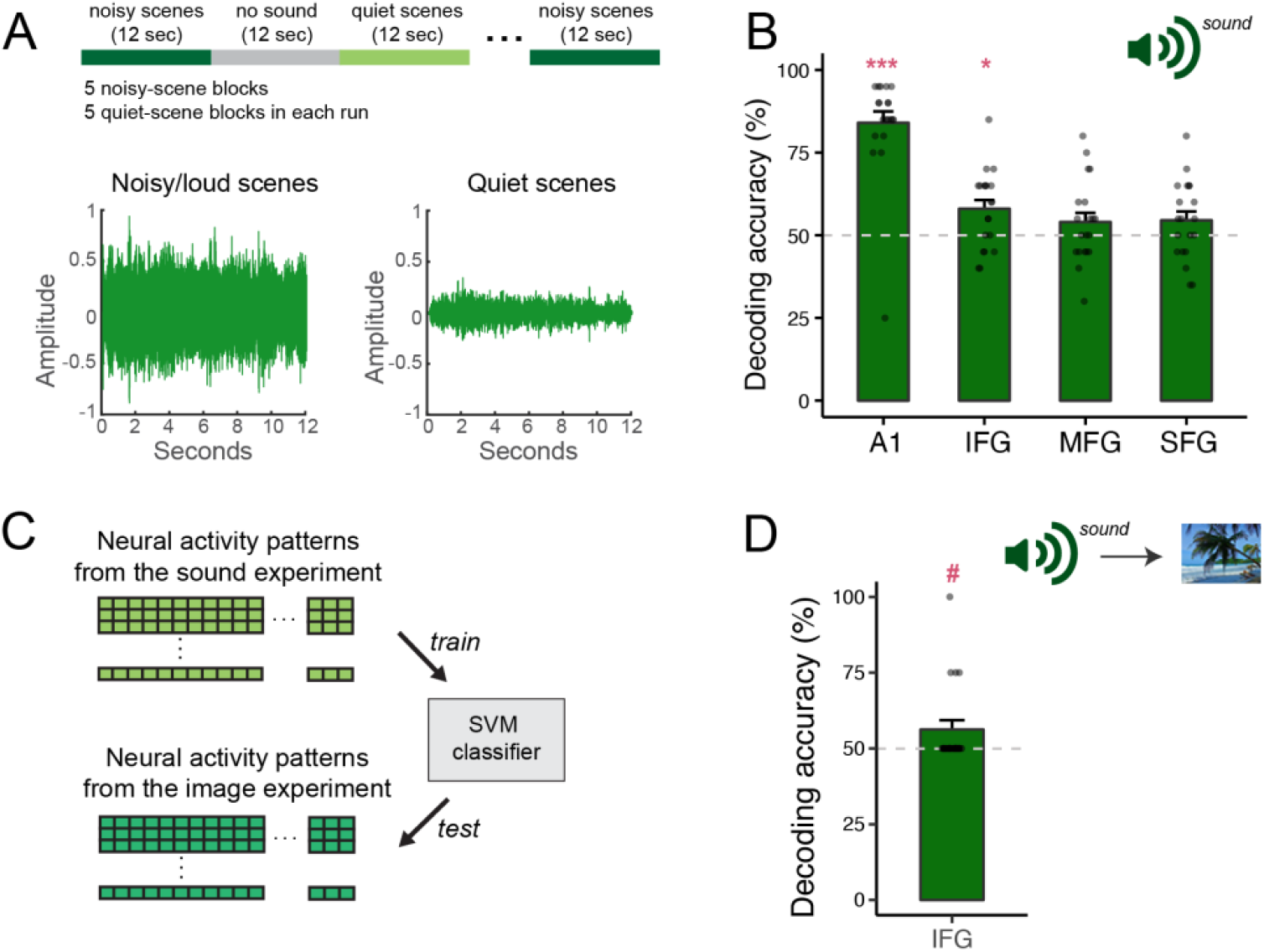
Decoding of sound properties. A) Illustration of the design of the sound experiment, and amplitude of an exemplar of the noisy sound clips and an exemplar of the quiet sound clips used in the localizer experiment. B) Sound-level decoding accuracy in each ROI C) Schematic of cross-decoding analysis across direct and indirect auditory percepts. D) Decoding performance of the cross-decoding analysis in IFG. Dots in each figure represent individual data points. #*p* < 0.05, ** *q* < 0.01, *** *q* < 0.001

As we confirmed that there are distinct patterns of neural representation for each type of sound level in both primary auditory cortex and PFC, we further examined whether these neural codes generalize to the sound level inferred from images. Cross-decoding of sound level was tested in IFG, which was the only brain area where we could decode both inferred (Figure 2B) and direct (Figure 4B) sensations of sound-level. In IFG, decoding succeeded at mean accuracy = 56.25%, *t*(19) = 2.032, *p* = 0.028 (Figure 4D).

These results indicate that in PFC, scene attributes are represented at an abstract level and generalize across sensory modalities, from direct sensations of temperature and sound level to scene properties inferred from visual percepts.

### Searchlight analysis

To explore how scene information is represented beyond the predefined ROIs, we performed a whole-brain searchlight analysis with a 5×5×5 voxels (10×10×10 mm) cube searchlight. The same decoding analyses for scene attributes (temperature and sound-level) and scene categories as in the ROI-based analysis were performed at each searchlight location using a linear SVM classifier, followed by a cluster-level correction for multiple comparisons.

Confirming the ROI-based analysis, both temperature and scene categories were decoded from PFC in the temperature judgment condition (Figure 5A). Clusters for category decoding were located in the anterior part of the MFG and SFG (highlighted in purple in Figure 5A), whereas the clusters for temperature decoding were located in posterior and medial part of PFC (highlighted in orange in Figure 5A) with little overlap.

**Figure 5.**
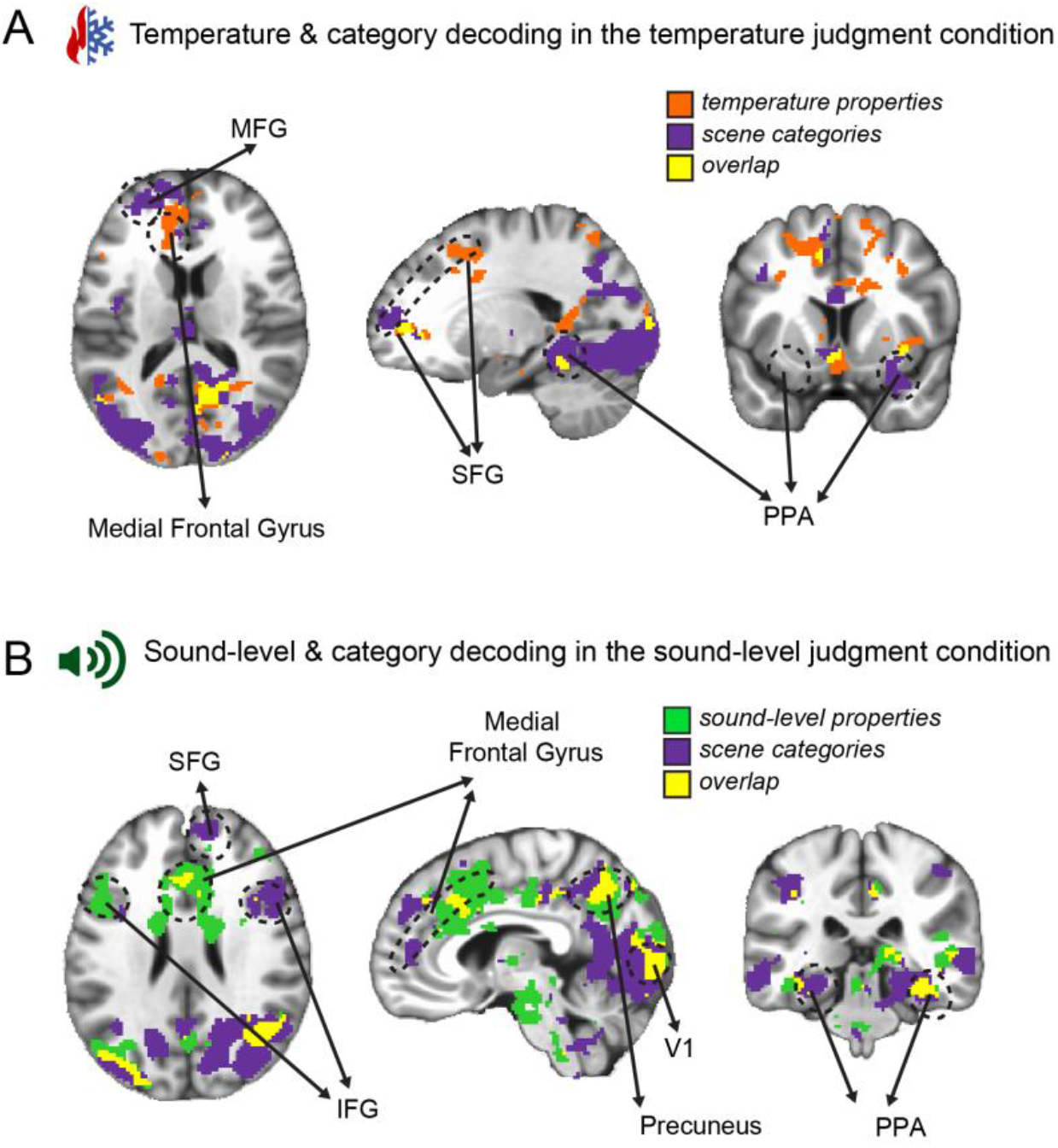
Searchlight maps of temperature and category decoding in the temperature judgment condition (A) and sound-level and category decoding in the sound-level judgment condition (B). Maps are thresholded at *p* < 0.05 and cluster-wise correction was conducted for multiple comparison (minimal cluster size of 215). MNI coordinate for (A): x= 20, y = −10, z = 15; for (B), x = −5, y = 32, z = 26.

In the sound-level judgment condition, scene categories were decoded in the anterior and posterior part of PFC, whereas the sound-level properties were decoded mainly in the medial prefrontal gyrus. A small overlap between the clusters can be observed in the MFG (Figure 5B).

Cross-decoding from the localizer experiment (direct percepts) to the main experiment (indirect percepts) was also performed in the whole brain using a searchlight analysis (Figure 6). Consistent to the ROI-based analysis, we identified clusters showing successful cross-decoding of temperature and sound level in the prefrontal regions. Specifically, for temperature decoding, we found three clusters overlapping with SFG in both hemispheres as well as with the right MFG, and the left IFG. For sound-level decoding, a cluster with successful decoding accuracy (*p* < 0.05) was located in the right MFG and IFG. We did not find any other clusters anywhere else in the brain that allowed for cross-modal decoding.

**Figure 6.**
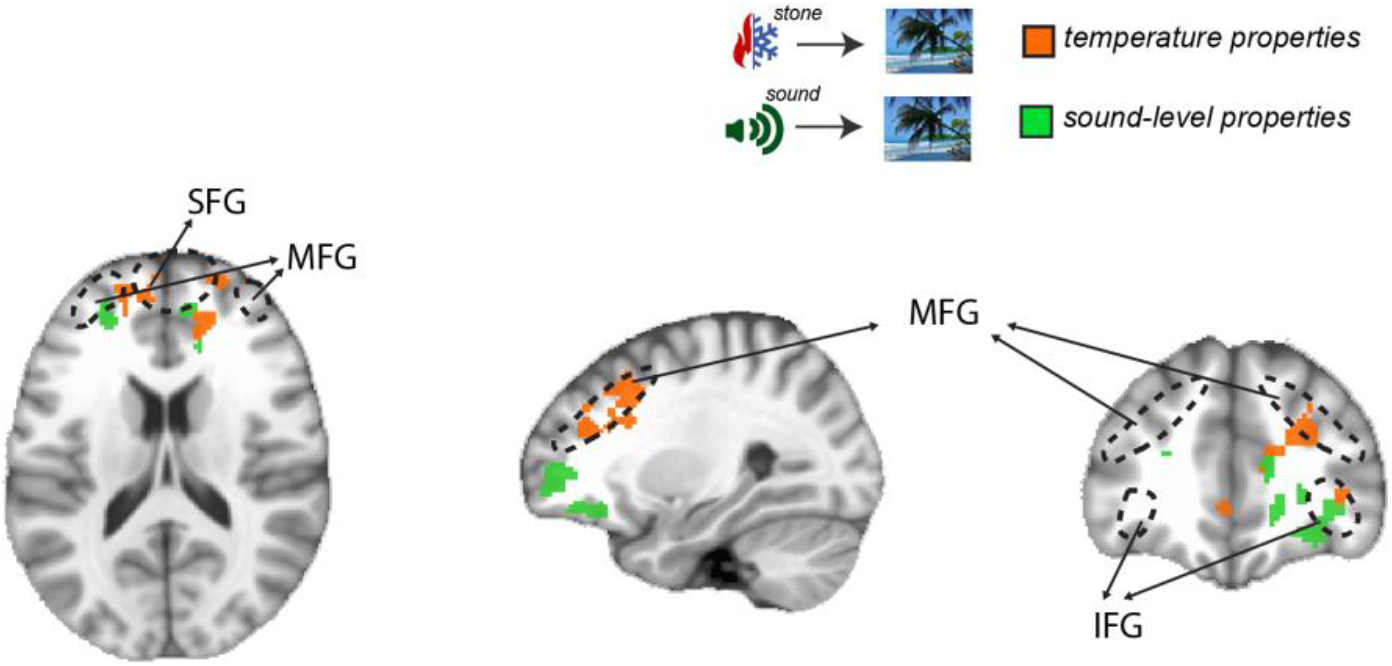
Searchlight maps of cross-decoding analysis for temperature (orange) and sound-level (green) decoding. Maps are thresholded at *p* < 0.05 and cluster-wise correction was conducted for multiple comparison (minimal cluster size of 215).

## Discussion

The present study demonstrates that both scene categories and scene attributes are represented in PFC. Specifically, scene attributes are represented in PFC only when the information is task-relevant. Across the two task conditions, the temperature of scenes was only decoded when participants were required to judge the temperature of a depicted scene, and the sound level was only decoded when participants were required to judge how noisy a scene was. Intriguingly, when scene properties were represented in PFC, they were computed at an abstract level, which transcended the level of specific sensory instances. Direct thermal or auditory percepts elicited similar neural activity patterns to those that were evoked when participants inferred these properties from scene images, suggesting that PFC generalizes across direct and indirect (inferred) perception.

The current study provides strong and clear evidence that PFC represents scene information, suggesting that scene information processing is not restricted to the scene-selective regions in the visual cortex but extends to the associative cortex. Our findings demonstrate that PFC represents both temperature and sound-level properties as well as scene category information. These findings are in line with previous studies that show that various aspects of scene information are processed beyond the scene-processing network in the visual cortex (Epstein & Baker, 2019; Groen et al., 2018; Jung et al., 2018).

The findings of the current study are also in line with previous work showing that neural representations in PFC can change according to task context (Bugatus et al., 2017; Harel et al., 2014; McKee et al., 2014). Both temperature and sound-level information were represented in PFC only when participants were making a decision about these properties; temperature properties were decoded in MFG when participants were judging scene temperature, but not when they were engaged in the sound-level judgment task. Similarly, sound-level properties of scenes were decoded in IFG and MFG only in the sound-level judgment condition. To some extent, these findings suggest that PFC might represent scene content that is not represented in the visual or auditory cortex; sound-level information of the scene was not represented in PPA or A1 but in PFC. Nonetheless, PFC represents this information when it is relevant to behavioral demands. These findings suggest that scene representations in PFC are largely modulated by task context or behavioral goals.

In contrast to scene attributes, the current study shows that scene categories are represented in both PPA and PFC regardless of task context. The decoding accuracy of scene categories does not vary systemically across different task conditions, suggesting that category information is represented in PFC even when category information is not directly related to behavioral demands. These findings are in line with previous behavioral studies, which showed that scene categories are processed automatically and obligatorily (Greene & Fei-Fei, 2014). Furthermore, several neuroimaging studies have shown successful decoding of scene categories using a passive-viewing paradigm without any task (Park et al., 2011; Walther et al., 2009, 2011). However, in these fMRI studies, participants could have been deliberately tracking categories of scenes, even though it was not explicitly required. The findings of the present study exclude this possibility by showing that categories are decoded even when the task manipulation is orthogonal to scene categories. This finding is limited by the use of only four scene categories in the current study. As there are hundreds of important scene categories (Tversky & Hemenway, 1983; Xiao et al., 2016), we cannot conclude that *all* of those categories are unaffected by task context. Future research is needed to further explore how tasks impact category representations in the brain.

Does the same set of voxels in the prefrontal cortex represent scene information, regardless of whether the scene information is about categories or scene attributes? Although our ROI-based analysis shows that the same regions (IFG and MFG) represent scene categories and scene attributes, the searchlight analysis indicates an interesting possibility that there may be different sets of clusters, each representing scene categories and scene attributes, respectively. Within the same anatomical ROIs, the clusters representing categories appear more anteriorly than the clusters representing scene attributes, which appear more posteriorly within the superior and middle prefrontal cortices (Figure 5A; the sagittal view). Further, in the sound judgment condition (Figure 5B), scene category representations appear in the right IFG, and representations of scene attributes appear in left IFG. Thus, it is possible that different parts of the prefrontal cortex each represent scene categories and scene attributes, thereby potentially explaining the different degree to which task influences these scene representations.

Our findings might appear to contradict previous work showing that distinct neural signals for categories are diluted in PFC (Bracci, Daniels, & Op de Beeck, 2017; Bugatus et al., 2017). For instance, Bugatus et al (2017) showed that the neural representations for visual categories, such as faces, words, and body parts, are evident in the ventral temporal cortex under different task contexts, but not in the prefrontal regions; PFC represents task context rather than categories of visual stimuli. There are, however, differences in the level of categories across the two studies (the current study and Bugatus et al., 2017) that could contribute to category-specific neural activity patterns in PFC. Unlike in Bugatus et al (2017), where superordinate categories, such as faces, pseudo-words, and body parts were used, the current study employs basic-level scene categories, which are the ‘default’ level of representing sensory input (Grill-Spector & Kanwisher, 2005; Jolicoeur et al., 1984; Mack et al., 2008). Several studies have demonstrated that such basic-level categories are accessible with very limited exposure (Mack et al., 2008; Mack & Palmeri, 2010) and with limited attention or cognitive resource (Greene & Fei-Fei, 2014; Li et al., 2002). Further, it should be noted that tasks are largely different across the two studies, and in the current study, the tasks in different conditions are still all related to scene understanding. Thus, future research is needed to better understand how different tasks may or may not impact category-selective representations in PFC.

Relatedly, Bracci et al., (2017) also show that neural representations of object categories are diluted by task context. PFC was found to represent object categories only when participants made judgments of semantic relationships among different objects (which referred as category task). The seeming inconsistency with our study might be due to different definitions of categories. We define scene categories as basic-level category (Tversky & Hemenway, 1983) and used multiple exemplars from each category (i.e., multiple beach images). Bracci et al., (2017) used different object categories under the same superordinate category (i.e., piano, violin, drum; for instruments) and defined category information based on semantic relationships among the individual items. Thus, although basic-level category information is preserved when the task is not related to category information (as found in the current study), richer information about the category, or semantic relationships among individual objects, might only be decodable from PFC when the task requires access to category information (as shown in Bracci et al., 2017).

Exploring this intriguing possibility in future work will require a different experiment design than is used in the current study.

The current study also broadens our understanding of what kinds of scene content are represented in the PPA. To our knowledge, our findings are the first to show that temperature can be decoded in PPA, just like scene categories (Walther et al., 2009) and scene properties related to spatial structure of a scene (e.g. openness, Park & Chun, 2009). The representation of temperature in the PPA appears to be driven by color to a large extent, suggesting that scene representations in PPA primarily rely on visual features.

We here provide evidence supporting the notion that representations in PFC generalize across sensory modalities, such as between the perceptions of images and stones, or images and sounds. Specifically, our findings are novel in that generalization across sensory modalities can occur when sensory information is inferred, not directly perceived. Although previous studies have shown that PFC represents sensory experience regardless of sensory modalities, participants in these studies always experienced direct stimulations in the corresponding modalities, such as hearing beach sounds and looking at a beach image (Jung et al., 2018; Vetter et al., 2014). The present study extends these findings by demonstrating that percepts inferred from a different modality (i.e., temperature or sound cues inferred from images) can elicit similar neural codes as the direct sensation would.

To summarize, the current study demonstrates that both scene categories and scene attributes are represented in PFC, however, neural representations of scene attributes (temperature and sound level) in PFC, not scene categories, are modulated by task context. These findings suggest that PFC flexibly represents scene content that is relevant to behavioral demands, allowing for efficient adaptation of behavior to varying situations in the real world.

## Conflict of Interest

The authors declare no competing interests

## Acknowledgement

The first author’s current affiliation is the following; Department of Psychology, Princeton University, NJ, USA. This work was supported by the Social Sciences and Humanities Research Council of Canada [430-2017-01189 to DBW], and the Natural Sciences and Engineering Research Council of Canada [RGPIN-2015-06696 to DBW].

## Author Contributions

YJ and DW designed the research, YJ performed research and analyzed the data under supervision of DW. YJ wrote the first draft of the paper, and YJ and DW edited the paper together.

## Supplementary materials

**Supplementary Figure 1.**
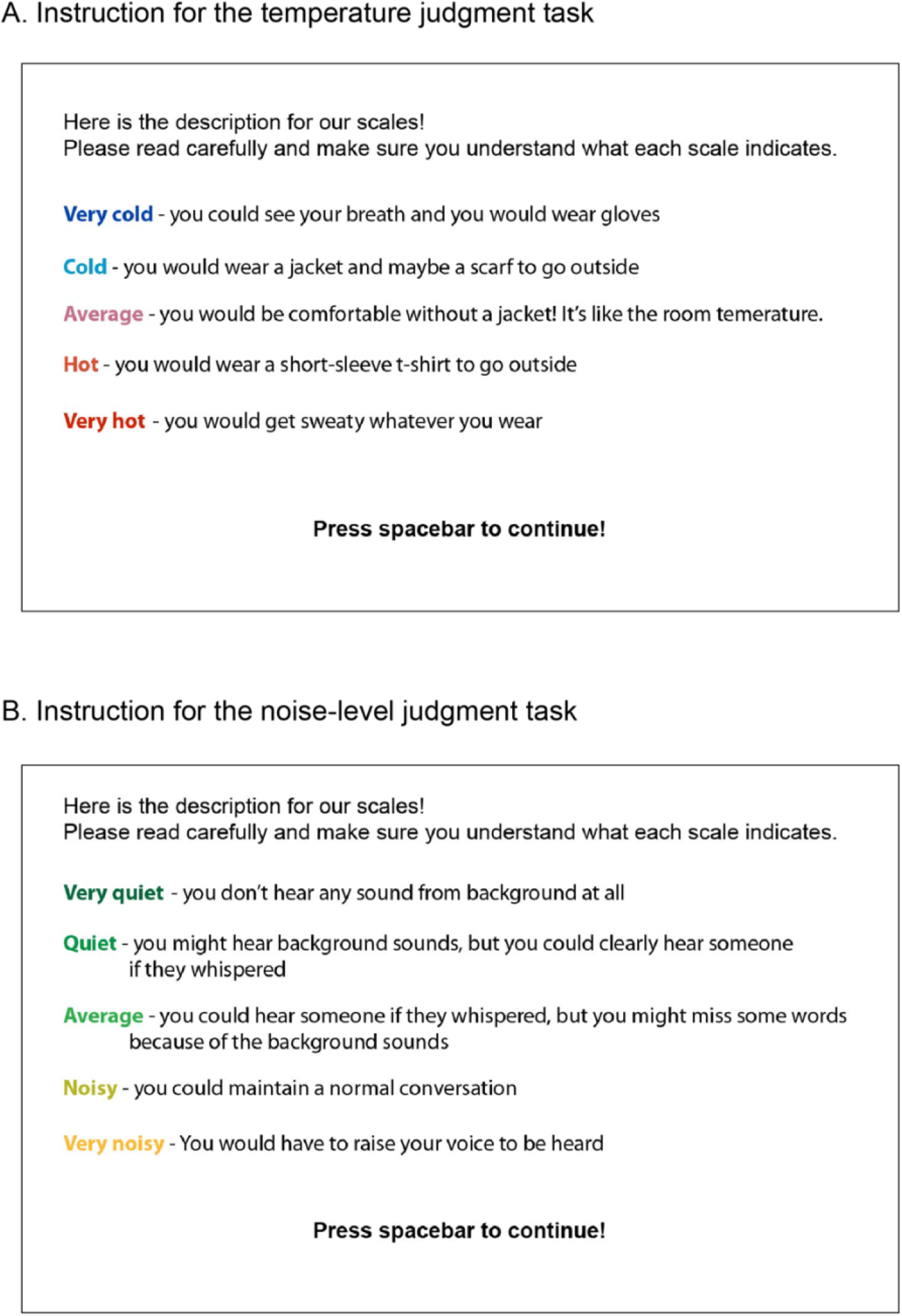
Instructions for the rating scales in the temperature judgment task (A), and in the sound-level judgment task (B).

**Supp. Figure 2.**
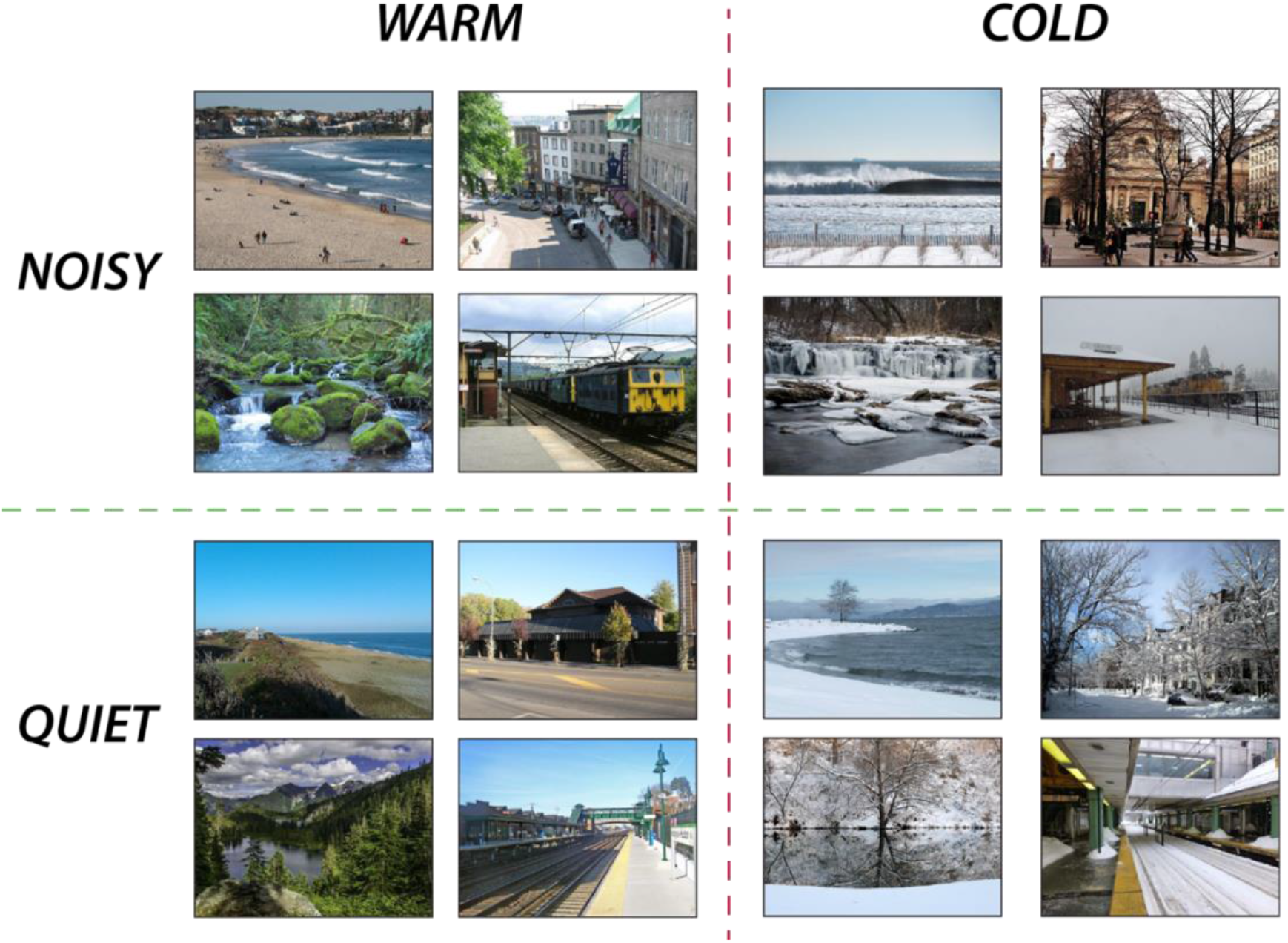
Examples of scene images used in the experiment. Based on the behavioral rating, each image was labeled as either warm or cold (temperature property) and at the same time, either noisy or quite (sound-level property). Also, these images belong to four basic-level scene categories: beach, city, forest, or train station. The entire stimuli set is available in our GitHub repository: (https://github.com/yaelanjung/scenes_fmri)

**Supp. Figure 3.**
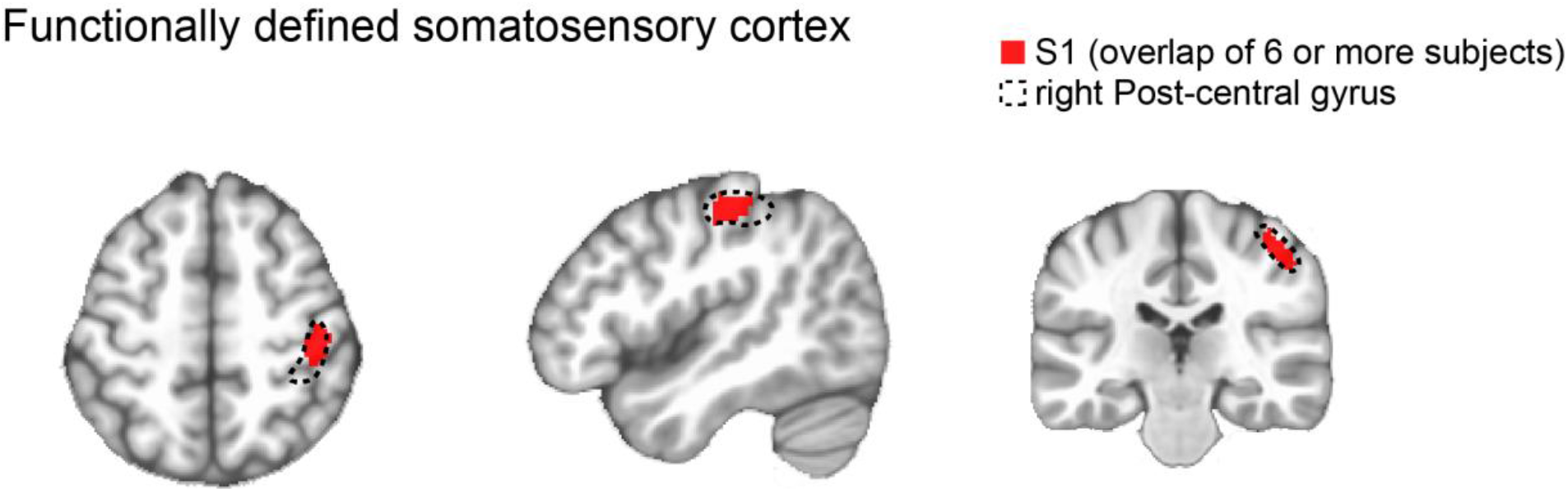
The localization of somatosensory cortex, where larger response was observed when participants were holding the stone in their left hand compared to when the stone stimuli was taken away (stone > no stone). The voxels where 6 or more participants show significant contrast (in stone > no stone) were first defined, and then this cluster (in red) was masked by the anatomical ROI of the right post-central gyrus (depicted in dotted line).

**Supp. Figure 4.**
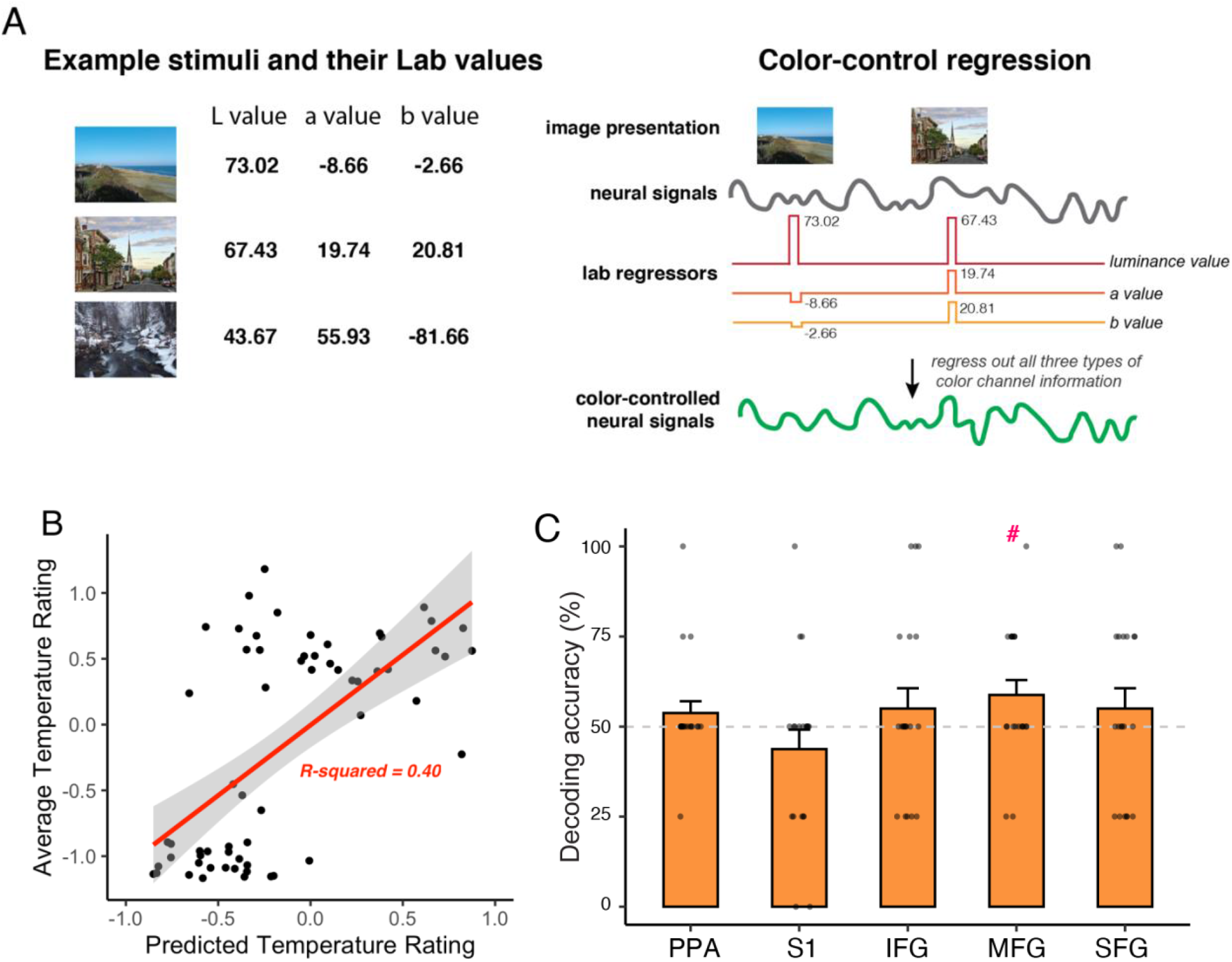
A. Example images and their average Lab values across all the pixels within each image. Each of these lab values were regressed out from the neural signal (see the left panel). B. Predicted temperature rating from all color-channel regressors (lab values) and the corresponding actual rating for individual images. C. Decoding accuracies of temperature (warm vs. cold) from images when LAB color values were regressed out from neural signals. #*p* < 0.05

**Supplementary Figure 5.**
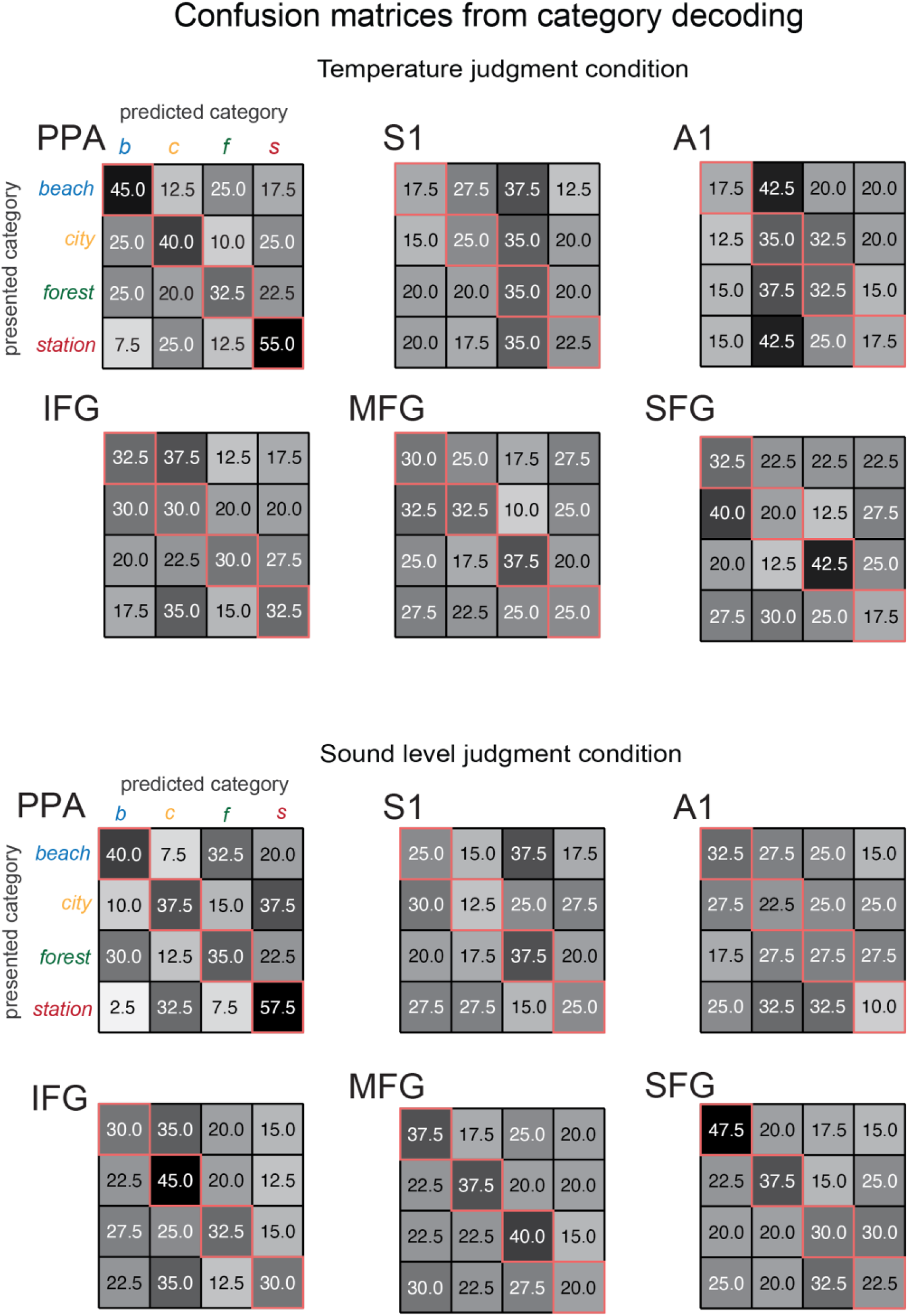

**Supplementary Figure 6.**
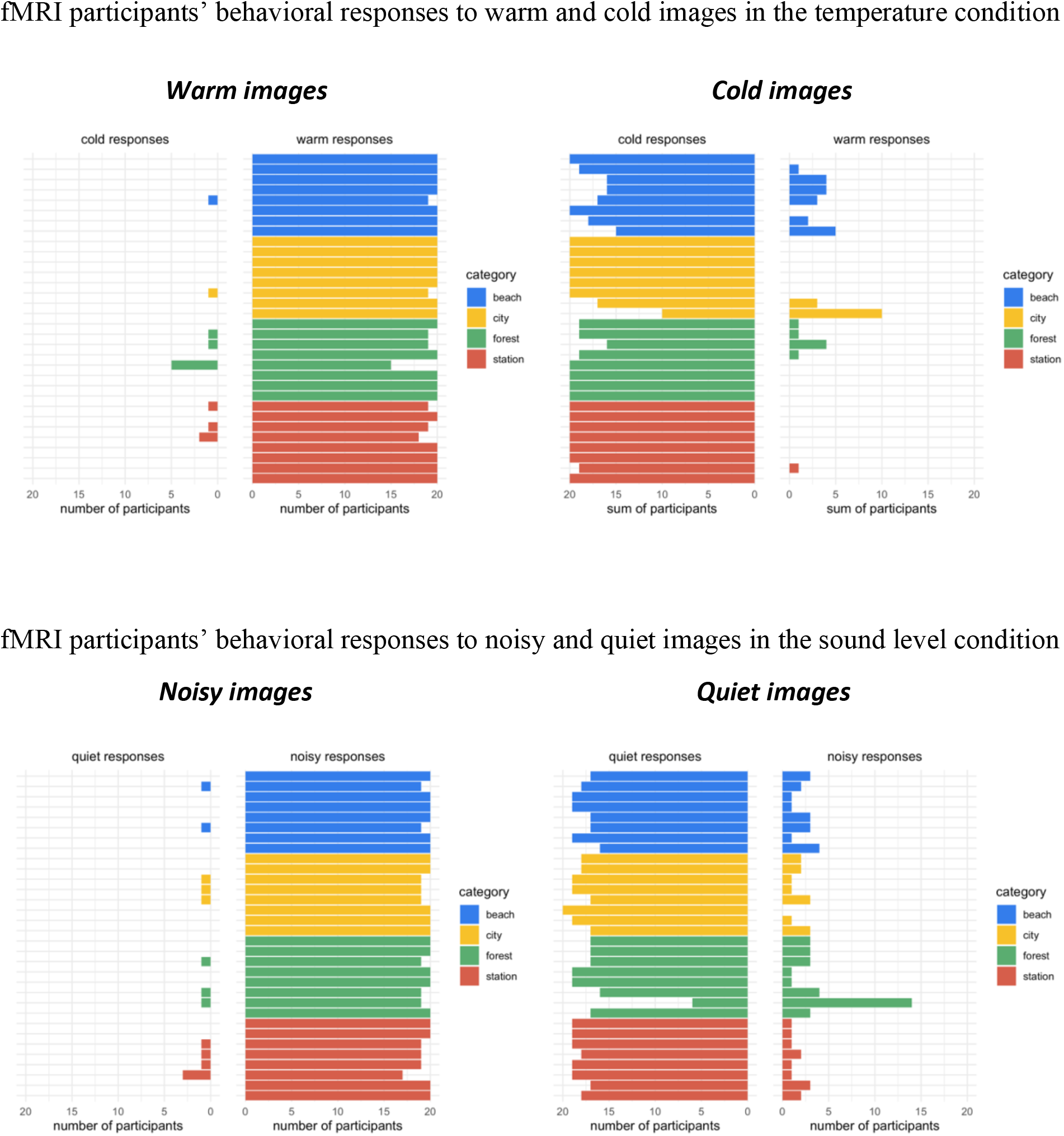

## Notes

### Competing Interest Statement

The authors have declared no competing interest.

